# MDR1 DNA glycosylase regulates the expression of genomically imprinted genes and helitrons

**DOI:** 10.1101/2024.07.31.606038

**Authors:** Kaitlin M Higgins, Jonathan Gent, Sarah N Anderson

## Abstract

Targeted demethylation by DNA glycosylases (DNGs) results in differential methylation between parental alleles in the endosperm, which drives imprinted expression. Here, we performed RNA sequencing on endosperm derived from DNG mutant *mdr1* and wild-type endosperm. Consistent with the role of DNA methylation in gene silencing, we find 96 gene and 86 TE differentially expressed (DE) transcripts that lost expression in the hypermethylated *mdr1* mutant. Compared with other endosperm transcripts, the *mdr1* targets are enriched for TEs (particularly Helitrons), and DE genes are depleted for both core genes and GO term assignments, suggesting that the majority of DE transcripts are TEs and pseudo-genes. By comparing DE genes to imprinting calls from prior studies, we find that the majority of DE genes have maternally biased expression, and approximately half of all maternally expressed genes (MEGs) are DE in this study. In contrast, no paternally expressed genes (PEGs) are DE. DNG-dependent imprinted genes are distinguished by maternal demethylation and expression primarily in the endosperm, so we also performed EM-seq on hybrids to identify maternal demethylation and utilized a W22 gene expression atlas to identify genes expressed primarily in the endosperm. Overall, approximately ⅔ of all MEGs show evidence of regulation by DNA glycosylases. Taken together, this study solidifies the role of MDR1 in the regulation of maternally expressed, imprinted genes and TEs and identifies subsets of genes with DNG-independent imprinting regulation.

**Significance Statement:** This work investigates the transcriptome changes resulting from the loss of function of DNA glycosylase MDR1, revealing that, in wild-type endosperm, targets of MDR1 are expressed predominantly from the maternal allele and this expression is suppressed in mutants. Furthermore, by combining expression data, DNA methylation data, and developmental expression data, we are able to categorize all maternally expressed, imprinted genes based on DNA glycosylase dependent or independent regulatory methods.

## Introduction/Background

DNA hypomethylation in the endosperm has been observed across flowering plants and results from the function of DNA glycosylases (Hsieh *et al*., 2009; Gehring *et al*., 2009; Zemach *et al*., 2010). DNA glycosylases remove DNA methylation through a base-excision repair-pathway whereby they remove 5-methyl-cystosine in the CG and CHG contexts preferentially targeting hemimethylated regions (Gehring *et al*., 2006), often nearby or in transposable elements (TEs) (Hsieh *et al*., 2009; Ibarra *et al*., 2012; Park *et al*., 2016). In Arabidopsis, the DNA glycosylase Demeter is responsible for the DNA hypomethylation observed in endosperm. Demeter functions in the central cell prior to fertilization, resulting in asymmetric hypomethylation specifically on the maternal allele (Ibarra *et al*., 2012; Park *et al*., 2016). In maize, two DNA glycosylases, MDR1 and DNG102, are expressed at a similar stage in development as Demeter. Maternal de-repression of R1 (*mdr1*) was first discovered due to its role in disrupting the characteristic expression of the gene red-color1 (R1) in maize kernels (Kermicle, 1995), Figure 1A/B). Recently MDR1 was mapped to a single gene and found to be a DNA glycosylase (Gent *et al*., 2022). MDR1, also known as ZmROS1b (Xu *et al*., 2022) or DNG101 (Jia *et al*., 2009), is active in the central cell and throughout endosperm development (Hoopes *et al*., 2019; Stelpflug *et al*., 2016). Prior research of *mdr1* mutants in maize revealed 18,464 DMRs between mutant and WT endosperm (Gent *et al*., 2022). Interestingly, double mutants of *mdr1*/*dng102* are not viable and cannot be passed through either male or female gametophyte. This suggests a critical role for DNA demethylation in seed development in maize.

**Figure 1:**
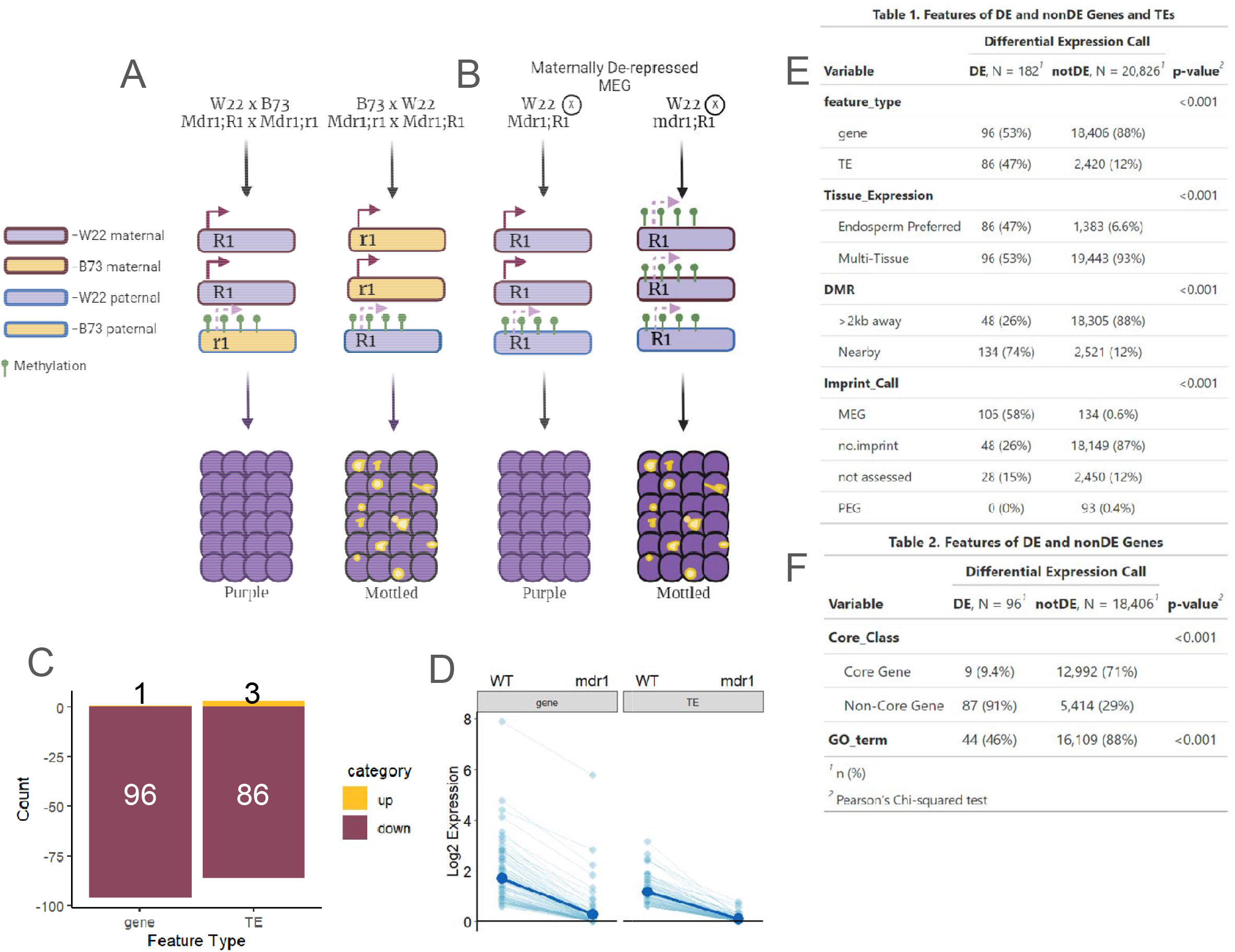
Genes and TEs are downregulated in mdr1 mutant endosperm in maize. Maternal De-Repression of R1 (MDR1) influences gene expression by demethylating and de-repressing gene expression in the endosperm. (A-B) Schematic of R1 expression in WT (A) and *mdr1* mutant (B) endosperm. (A) In a reciprocal cross between W22 with the dominant red-color1 (R1) allele and non-pigmented B73 with the recessive r1 allele, the kernels display a solid purple color when R1 is inherited maternally due to low maternal DNA methylation, and mottling when R1 is inherited paternally due to high paternal DNA methylation. (B) When W22 (R1;MDR1) is self pollinated the kernels are solid purple colored, however in the *mdr1* mutant, the self-pollinated kernels are mottled due to high DNA methylation of the maternal alleles. (C) Differentially expressed genes and transposable elements (TEs) in *mdr1*. DE genes used for downstream analyses include 96 genes and 86 TEs that are down in mutants relative to WT. (D) Magnitude of expression change in *mdr1* mutants. Data show relative expression (log2(mean rpm +1)) for the mean of expression in WT and mutant for DE genes (left) and TEs (right). Individual features are connected by lines. Black line at bottom denotes a value of zero. (E-F) Table of features of DE and nonDE genes and TEs in this study. Tables list the number and percent of features in each intersect, along with the p-value from Pearson’s chi-squared test to determine significant differences between DE and nonDE features. (E) DE genes and TEs are defined as endosperm preferred when more than 65% of the sum of expression across tissues comes from endosperm replicates. DMRs are classified as Nearby when the WT vs *mdr1* mutant DMRs overlap or are within 2kb of features. Imprinting calls were taken from Anderson et al. 2021 and genes are classified as MEGs or PEGs when called in either crosses between W22 and B73 or Ph207. (F) Core gene classifications are based on presence across all 26 NAM genomes in Hufford et al. 2023. Genes are further classified based on whether or not that have at least one Gene Ontology (GO) annotation assigned.

One major consequence of asymmetric DNA hypomethylation in the endosperm is genomic imprinting (Figure 1A). Imprinting is parent-of-origin biased gene expression and can present as either Maternally Expressed Genes (MEGs) or Paternally Expressed Genes (PEGs). Misregulation of specific imprinted genes leads to seed size distortion and often inviability (Tiwari *et al*., 2010; Hornslien *et al*., 2019; Yuan *et al*., 2017). Imprinting has been observed and studied in a variety of plant species, however the mechanisms underlying imprinting regulation are not fully understood. The best understood model of regulation exists for maternally activated MEGs, which are targets of DNA glycosylases in the central cell and are primarily expressed from the maternal allele in the endosperm (Zhang *et al*., 2014; Batista and Köhler, 2020). However, this subset of MEGs has not yet been well defined in maize. Furthermore, DNA glycosylases have also been predicted to result in the expression of some PEGs through demethylation of respressors (Anderson and Springer, 04/2018), although the extent to which this occurs *in vivo* is unknown.

Here, we perform RNA sequencing of *mdr1* mutant endosperm to reveal genes and TEs whose expression is regulated by DNA glycosylase activity. We find nearly 200 genes and TEs that display lower expression in the mutant relative to wild-type, consistent with the silencing role of DNA methylation. DE features (genes and TEs) share several key features, including an enrichment for expression primarily in the endosperm, and an enrichment for nearby DMRs between endosperm and embryo. Additionally, the majority of differentially expressed (DE) genes are maternally expressed (MEGs) and show expression patterns consistent with the maternal derepression model of imprinting. However, approximately half of all MEGs are DE in *mdr1* mutants. By utilizing multi-omic integration of expression in mutants and across development, DNA methylation in inbred and hybrid mutants, and previously published imprinting data, we were able to quantify the extent of imprinting resulting from maternal derepression.

## Results

To test how *mdr1* affects transcription within the endosperm, we performed RNA-seq on endosperm from WT W22 and *mdr1* mutant W22 endosperm at 14 days after pollination. Reads were mapped to the W22 genome (Springer *et al*., 2018) and reads were assigned to both genes and TEs as in (Anderson *et al*., 2019). We then analyzed differential expression of uniquely mapping genes between WT W22 and *mdr1* W22 and found a total of 97 genes and 89 TEs that are DE between the mutant and WT. We found an overwhelming bias towards loss of expression in the mutant, with only 1 gene and 3 TEs upregulated (Figure 1C). This result is consistent with the canonical knowledge of DNA methylation in reducing gene expression and the role of MDR1 in demethylation in WT endosperm (Figure 1A). Given the primary role of MDR1 in gene activation and the observation that the single upregulated gene (Zm00004b027381) has no known function, we focused downstream analyses on only the genes with repressed expression in the mutant. This resulted in 96 genes and 86 TEs in our DE feature set, all of which are expressed lower in the hypermethylated *mdr1* endosperm relative to WT (Figure 1D). When compared with expressed but nonDE features, DE features are enriched for TEs, with 47% of DE features annotated as TEs, compared with only 12% for expressed but nonDE features (Figure 1E).

We next sought to find defining features of DE genes and TEs. Many of the genes in our DE list lacked functional information. When we looked for GO terms in our DE genes, we found 44 (46%) had associated terms; in contrast 16,109 (88%) of nonDE genes had associated GO terms (Figure 1F). We further assessed the conservation of our DE genes within maize. To assess conservation within maize, we determined how many of these DE genes are considered “Core Genes’’ within the 26 founder genomes of the maize Nested Association Mapping (NAM) genomes which have been fully assembled and annotated (Hufford *et al*., 2021). In B73, 27,910 genes (58% of all annotated genes) are considered core genes. To identify W22 syntelogs of core genes, we compared single copy genes between B73 and W22. Of these, 13,781 B73 identifed core genes are also present in W22, yet only 9 core genes (9.3% of DE) are differentially expressed in *mdr1* mutant endosperm (Fig 1F). The relatively small number of genes conserved across maize suggests that genes with expression controlled by MDR1 are primarily dispensable genes without functional annotations.

To further understand the broader expression context of DE genes, we next aimed to assess the developmental expression patterns of genes in W22. To accomplish this goal, we used a W22 expression atlas with two biological replicates of samples from 10 tissues (endosperm, embryo, anther, tassel, immature ear, leaf, 10th leaf, internode, root, and shoot and coleoptile) and mapped to the W22 genome (Monnahan et al. 2020). To simplify our analysis, we categorized expression as endosperm preferred, defined as more than 65% of normalized transcripts across all tissues originating in the two endosperm replicates, or multi-tissue, defined as less than 65% of all transcripts were originating in the endosperm replicates. Genes and TEs that are expressed in the endosperm, but are nonDE in *mdr1*, show a wide variety of expression patterns across tissues. In contrast, DE genes show a much more limited range of expression with expression primarily restricted to the endosperm (Figure S1). To quantify this observation, we assessed the percentages of DE and nonDE genes and TEs that fall into each category of expression. This revealed that 71% of DE and 6.7% of non-DE genes were defined as endosperm preferred, representing a significant enrichment for endosperm preferred expression for DE genes and TEs (Figure 1E). This enrichment of endosperm-preferred expression is consistent with MDR1’s expression primarily in the endosperm and its functions in gene activation.Taken together these results show that DE genes that are repressed in *mdr1* mutants are often associated with a lack of functional annotations, a lack of conservation within maize and with sorghum, and wild-type expression primarily restricted to the endosperm.

### MDR1 targets are demethylated in WT endosperm

In our previous study characterizing the *mdr1* mutant, we assessed DNA methylation between *mdr1* and WT endosperm to identify Differentially Methylated Regions (*mdr1*-DMRs). The majority of DMRs had higher CG and CHG methylation in *mdr1* consistent with MDR1’s predicted 5-methylcytosine glycosidic function, so the *mdr1*-DMR set is composed of 18,464 DMRs with higher methylation in the mutant (Gent *et al*., 2022). To identify cases where MDR1’s demethylase activity is associated with the differential expression of the genes, we evaluated the location of the closest *mdr1*-DMRs to DE features. For 62.5% of DE genes and 67.4% of DE TEs, DMRs overlap the gene or TE annotation, compared with only 2.2% and 7.5% of nonDE genes and TEs, respectively (Figure 2A). However, regulatory regions are often nearby rather than directly overlapping genes, so we categorized genes as nearby *mdr1*-DMRs when they overlap or are within 2 kb of an *mdr1*-DMR. Among DE genes and TEs 74% (134) had nearby *mdr1-*DMRs, whereas just 12% of nonDE genes and TEs had nearby *mdr1-*DMRs. This revealed that DE genes and TEs are 28-fold enriched for having a nearby *mdr1*-DMR compared with nonDE genes and TEs (p < 2.2 e-16, chi square test). To quantify where in DE genes DMRs are most often observed, we analyzed the occurrence of DMRs at the 2 kb centered around the TSS and poly-A sites for DE and nonDE genes (Figure 2B). This revealed that DE genes have a DMR near their TSS 44-times more often than expected based on the distribution of control regions, whereas expressed but nonDE genes were enriched only 2-fold over control regions (Figure 2B).

**Figure 2:**
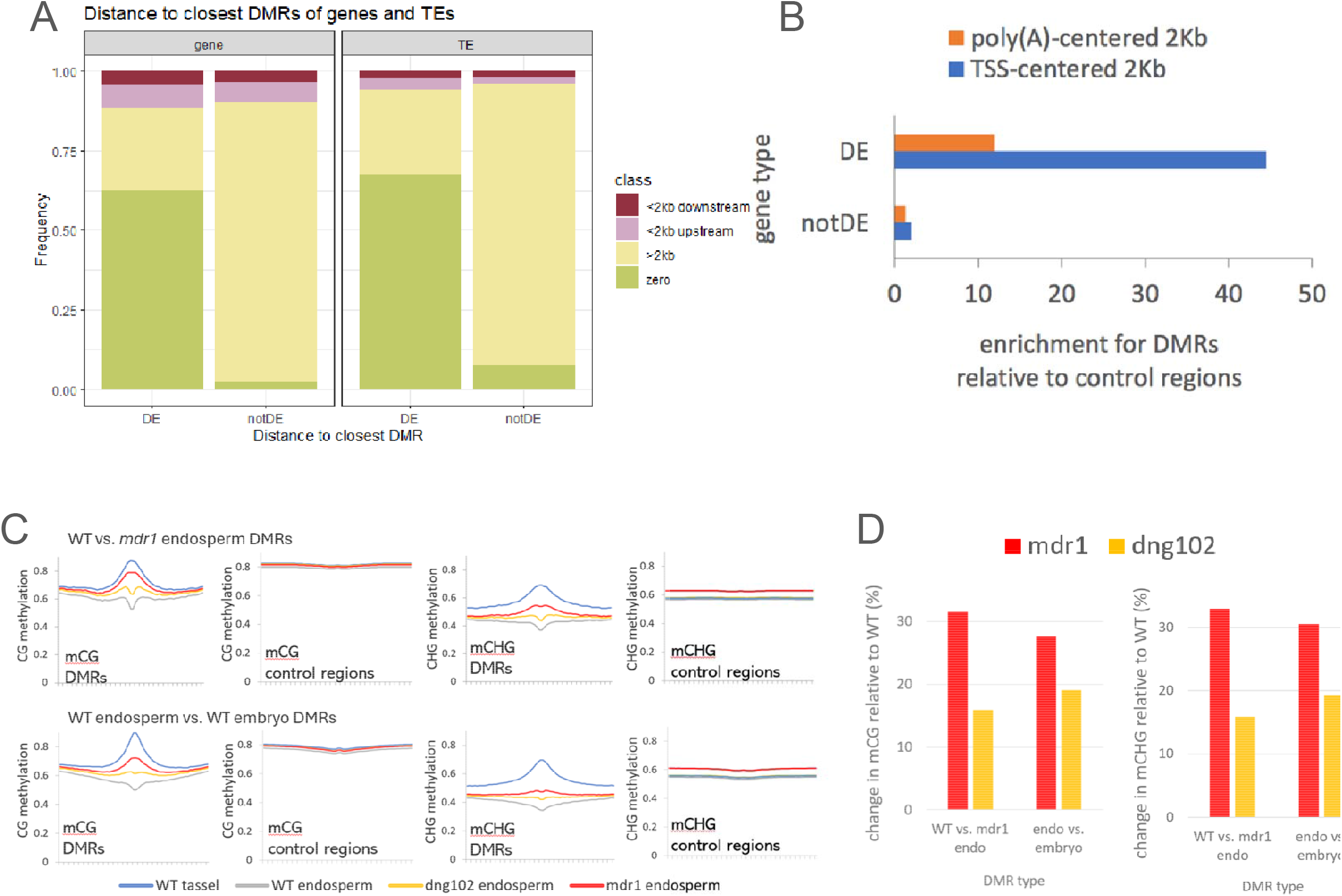
DE genes are differentially methylated in the mutant. Differentially Methylated Regions (DMRs) between WT and *mdr1* mutant endosperm are enriched near DE genes and TEs. (A) Distance to nearest DMR for DE and nonDE genes and TEs. DE genes and TEs are enriched (p <2.2 e-16, chi-square test) for having overlapping or nearby DMRs when compared to nonDE genes and TEs. (B)Enrichment of DMRs overlapping the 2kb regions centered on transcription start sites or polyadenylation sites for DE and nonDE genes using control regions (200-bp regions that were eligible for identifying DMRs based on read coverage and number of informative cytosines) throughout the genome as a baseline for enrichment comparison. (C) DNA methylation metaplots in the CG and CHG contexts for Endosperm vs Embryo DMRs (top) or *mdr1* endosperm vs WT endosperm DMRs (bottom) compared to control regions. Lines represent methylation in WT tassel (blue), WT endosperm (gray), dng102 endosperm (yellow), and *mdr1* endosperm (red). (D) Summary of DNA methylation change in *mdr1-*DMRs and endosperm vs embryo DMRs in the CG(left) and CHG(right) contexts in *mdr1* (red) and *dng102* (yellow)

While MDR1 is responsible for many sites of demethylation in the endosperm, it has a paralog (DNG102) that is also active within the endosperm, and at least one functional copy of at least one of these genes is required for gametophyte formation (Gent *et al*., 2022). Given the overlap in function, we sought to characterize the methylation levels in the *dng102* single mutant to understand if these genes act in overlapping or independent regions of the genome. Mature endosperm was harvested from *dng102* mutant endosperm and assessed for CG and CHG methylation over *mdr1*-DMRs and control regions. We found that *mdr1*-DMRs are also hypermethylated in *dng102* mutants though to a lesser extent (Figure 2C). To quantify this, we evaluated the total methylation change in two DMR sets: *mdr1*-DMRs and wt-DMRs (Figure 2D). The wt-DMR set includes 52,919 DMRs that are hypomethylated in the WT endosperm when compared with the WT embryo. When directly comparing the methylation changes above *mdr1-*defined DMRs, *dng102* causes around half of the total change in methylation that *mdr1* does, whereas in the endosperm vs embryo DMRs the total change is more similar between the two mutants (Figure 2D), though the change in *dng102* is still less drastic than in *mdr1*. This indicates that the two paralogs display functional redundancy at some loci.

### MDR1 DE targets and the association with transposable elements

Transposable elements contribute a major portion of the maize genome and are generally methylated and transcriptionally silent. However, TEs can be a source of regulatory elements that are variable across genotypes and across tissues (Makarevitch *et al*., 2015; Warman *et al*., 2020; Anderson *et al*., 12/2019; Liang *et al*., 2020). Prior work has linked MDR1 to demethylation of transposable elements (Xu *et al*., 2022) as well as hypothesized that imprinting arises as a byproduct of demethylase activity on TEs (Hsieh *et al*., 2009; Gehring *et al*., 2009; McDonald *et al*., 2005). Among the 2509 TEs transcribed in the endosperm, 3.4% are DE, compared with 0.5% for genes, suggesting that TEs are over-represented in our DE feature set relative to genes. In addition to expression changes in TEs themselves, methylation and expression changes in TEs may also contribute to expression changes in nearby genes.

To delve into the extent of TE influence on DE genes in *mdr1*, we evaluated both TEs that were differentially expressed (DE TEs) as well as the TEs closest to DE genes (close TEs) for overrepresented superfamilies and families. Of the 89 DE TEs, 40 (45%) belonged to the helitron (DHH) superfamily, compared with 1.6% for expressed non-DE TEs (Figure 3A). Of the close TEs, we discovered a significant enrichment of helitrons being the closest TE to DE genes (Figure 3B, p = 6.02e-11, hypergeometric test). Although there are 40 helitrons that are differentially expressed and 40 helitrons that are closest to differentially expressed genes, only 7 are both DE and the closest to a DE gene. This shows that these enrichments are not due to complete overlap of DE TEs and close TEs, but rather independent enrichments.

**Figure 3:**
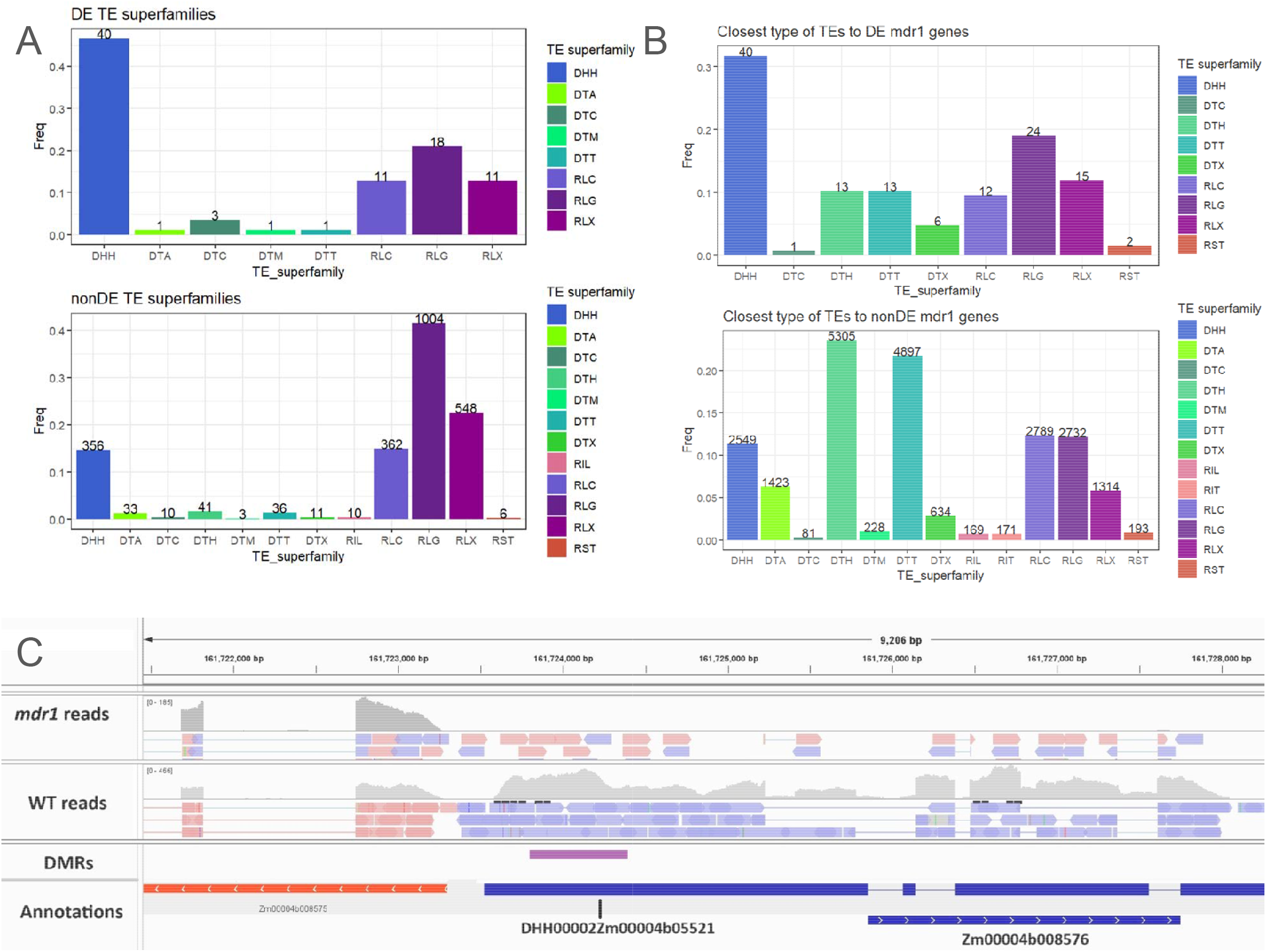
Transposable Elements and their association with DE *mdr1* features. (A) DE TEs and TE relationships to DE genes Breakdown of the TE superfamilies of DE (top) and nonDE (bottom) TEs. Total count of TEs is listed above each bar. (B) Closest superfamily of TE to DE (top) and nonDE (bottom) genes. Total count of TEs is listed above each bar. (C) IGV visualization of read alignment of overlapping DE TE-gene pair Zm00004b008576 and DHH00002Zm00004b05521. Gray histograms show the coverage of RNA-seq reads, and individual reads are colored based on forward (blue) or reverse (red) strand mapping.

We next evaluated the specific families of DE and close TEs. Of the DE helitrons, 32 belonged specifically to the DHH00002 family. This represents a 4.8-fold enrichment of this family of TEs in the DE set relative to the non-DE set (p < 7e-17, hypergeometric test). Further, when we explored the specific families present in the subset of helitrons nearby DE genes we found a significant enrichment of the DHH00002 helitron family (p = 2.21e-12, hypergeometric test). DHH00002 was previously shown to exhibit an enrichment for imprinted expression, with a total of 6 DHH2 TEs both repressed in *mdr1* mutants and maternally expressed in crosses between B73, W22, and PH207 (Anderson *et al*., 2021). Taken together these results make the DHH00002 family transposons a likely candidate for targeting by MDR1.

Finally, we evaluated whether any of the DE TEs overlap DE genes, and found 9 overlapping DE gene/TE pairs. Of these, 6 genes are fully within TEs and the other 3 have partial overlaps between the gene and TE annotations (Table S1). By visualizing read alignments to these gene/TE pairs, we find that RNA-seq reads span the length of both the gene and TE annotations (Figure 3C, Figure S2), suggesting that these co-occurring DE calls may result from fused transcripts rather than transcription from gene and TE features independently.

### MDR1 targets have maternally biased expression in WT endosperm

Since the *mdr1* mutation was discovered for its role in maize imprinted expression, we assessed the overlap between DE genes and previously determined imprinting calls for W22 genes in reciprocal crosses with B73 and Ph207 (Anderson *et al*., 2021). We found that DE genes are enriched for maternal expression (p=7.89 e-61 hypergeometric test) with 106 of 182 DE features (58%, including 57 genes and 49 TEs) classified as MEGs or matTEs. In contrast, only 0.4% (82) of non-DE genes and 2% (52) of non-DE TEs are maternally expressed genes (MEGs) or maternally expressed TEs (matTEs). Interestingly, DE MEGs/matTEs and DE genes that are not imprinted have a similar rate of overlap with *mdr1*-DMRs, while non-DE MEGs/matTEs overlap *mdr1*-DMRs far more often than genes that are neither DE nor imprinted (Figure S3A). To understand this observation further, we assessed the parental bias in gene expression for DE and non-DE genes in reciprocal crosses between W22 and either B73 or Ph207 from our previous study of imprinting in maize (Anderson et al., 2021). Plotting parental bias for DE and non-DE genes reveals that, while non-DE genes have a distribution centered near biparental expression (RER = .66), the mean of parental bias (RER) for DE genes is .94, which indicates a substantial bias towards maternal expression (Figure 4A, Figure S3B). This observation suggests that although 59% of DE genes and 57% of DE TEs are MEGs or matTEs (Figure 4B), DE genes generally exhibit maternally biased expression regardless of whether or not they met the stringent threshold to be classified as MEGs/matTEs previously.

**Figure 4:**
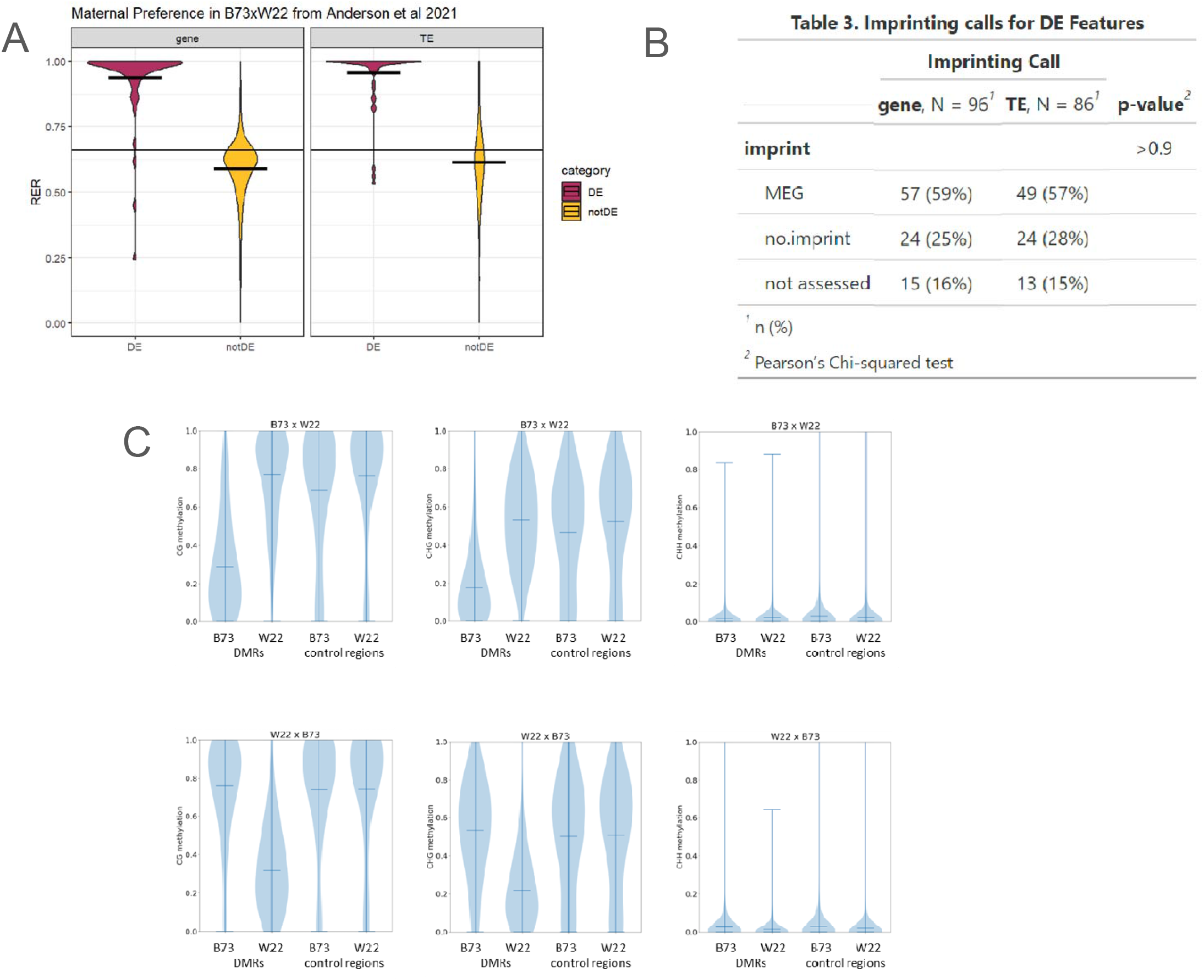
DE genes and imprinting. MDR1 targets the maternal alleles of imprinted genes. (A) Violin plot of parent-of-origin biased expression for DE vs. nonDE genes (left) and TEs (right). Data is from Anderson et al 2021 and shows Reciprocal Expression Ratio (RER) a metric calculated by first mapping allele-specific reads from reciprocal crosses to both parental genomes and then dividing the normalized reads for each allele when inherited maternally by the sum of maternal and paternal expression. This results in a value ranging from 0 to 1 where 1 results from exclusively maternal expression, 0 results from exclusively paternal expression, and 0.66 results from equal expression from the two maternal and one paternal allele in the endosperm (horizontal line). Horizontal lines crossing the violin plots indicate the mean value for each set. (B) Table of imprinting calls for DE genes and TEs. (C)Parent-specific DNA methylation in hybrid endosperm. Violin plots showing CG (left), CHG (middle), and CHH (right) methylation for B73 and W22 alleles for DMR (embryo vs endosperm) and control regions (all regions with sufficient EM-seq coverage) in mature endosperm of B73 x W22 (top) and W22 x B73 (bottom) crosses, where the maternal genome is listed first. Horizontal lines indicate mean values..

Prior studies on demethylation by DNA glycosylases (DNGs) in endosperm have focused on identification of regions that are differentially methylated between parental genomes. Even in maize, where DNGs are are strongly expressed after merging of the two parental genomes in the same nucleoplasm in endosperm, it is clear that differential methylation between parental genomes is due to demethylation of the maternal genome (Zhang *et al*., 2014). These results, however, do not rule out the possibility that DNGs can also demethylate both genomes. Regions where both genomes are demethylated would not have been identified in studies that focus on differential methylation between parental genomes. To address this possibility, we generated DNA methylomes from W22 X B73 and B73 x W22 hybrid endosperm from dry seed and generated genome-specific methylation calls. EM-seq reads that overlap sequence variants allow for assigning to their source parental genomes in a hybrid and allow for assigning genome-specific methylation calls to all cytosines spanned by the reads. Thus in a single hybrid endosperm, we could generate methylation calls for each genome at loci with uniquely mappable reads, where “uniquely mappable” means not just a single-copy sequence in the W22 genome, but also one that has a variant that allows it to be distinguished from the homologous B73 locus. We then used these parent-specific methylation values to measure methylation of previously identified regions that are demethylated in endosperm relative to embryo (Gent *et al*., 2022). We found no evidence for demethylation of the paternal genomes at these loci for either hybrid (Figure 4D). These results indicate that MDR1 and DNG102 are specifically targeted to the maternal genome at these loci, either by limiting their activity to before karyogamy or by an unidentified epigenetic distinction between the parental genomes that carries over into the endosperm. MDR1 acting specifically on the maternal allele, along with the differential expression of 106 genes and TEs previously identified as imprinted suggests MDR1 alone is responsible for imprinted expression of nearly half of maternally expressed genes in W22.

### Only a subset of MEGs are regulated by maternal de-repression

While MEGs/matTEs are clearly enriched among DE genes, less than half of all maternally expressed genes and TEs are DE in *mdr1* mutant endosperm. There are a few possible explanations for this observation. First, some MEGs/matTEs are likely redundantly regulated by glycosylases MDR1 and DNG102, resulting in expression that is not affected in the single MDR1 mutant. Alternatively, other non-DE MEGs/matTEs may result from imprinting regulation that is not dependent on the DNGs. Prior studies of imprinting in maize have defined two categories of MEGs, maternally de-repressed MEGs and paternally repressed MEGs (Zhang *et al*., 2014). Maternally de-repressed MEGs will typically exhibit expression specifically in the endosperm (Zhang *et al*., 2014; Batista and Köhler, 2020). While MDR1-dependent MEGs are all by definition maternally de-repressed, we expect the nonDE MEGs to contain both redundantly regulated maternally de-repressed MEGs displaying endosperm specific expression as well as having DMRs nearby, and paternally repressed MEGs that are expressed in various tissues and do not have nearby DMRs. To capture more of the differential methylation that would contribute to maternal de-repression of MEGs, here we switch to utilizing the wt-DMRs which contain DMRs between embryo and endosperm and are not limited by redundancy between MDR1 and DNG102. We tested our hypothesis by comparing expression patterns across development as well as distance to nearest wt-DMR for DE and nonDE MEGs/matTEs.

First we looked at expression for DE and nonDE MEGs/matTEs across tissues to determine if there were major differences in the tissue specificity of expression. We found that while a portion of each feature set displayed endosperm preferred expression, the nonDE MEGs/matTEs exhibit a clear split between endosperm preferred and multi-tissue expression (Figure 5A). While the majority of DE MEGs/matTEs are endosperm preferred (81%), only 46% of nonDE MEGs/matTEs show this expression pattern (Figure 5B). Next we analyzed the distance to the closest wt-DMR for each MEG/matTE. This revealed that while the majority (89%) of DE MEGs/matTEs are near wt-DMRs, fewer (77%) of nonDE MEGs/matTEs are near wt-DMRs (Figure 5B). Combining these two analyses, we then evaluated how closely the DE and nonDE MEGs/matTEs follow the two models for maternal expression by looking at the proportions of each category that contain a nearby wt-DMR and display endosperm preferred expression (Figure 5C). This revealed that 77 (72%) of DE MEGs/matTEs are both endosperm preferred as well as have a nearby wt-DMR as expected for maternally de-repressed MEGs/matTEs. In contrast, only 39% of nonDE MEGs/matTEs were both endosperm preferred and have a nearby wt-DMR. Additionally, more than 15% of nonDE MEGs/matTEs show no features of maternal de-repression at all. This suggests that while MEGs/matTEs that are repressed in *mdr1* mutant are clear examples of maternal de-repression by *mdr1*, the nonDE MEGs/matTEs likely consist of a mix of redundantly regulated transcripts and unrelated imprinting control.

**Figure 5:**
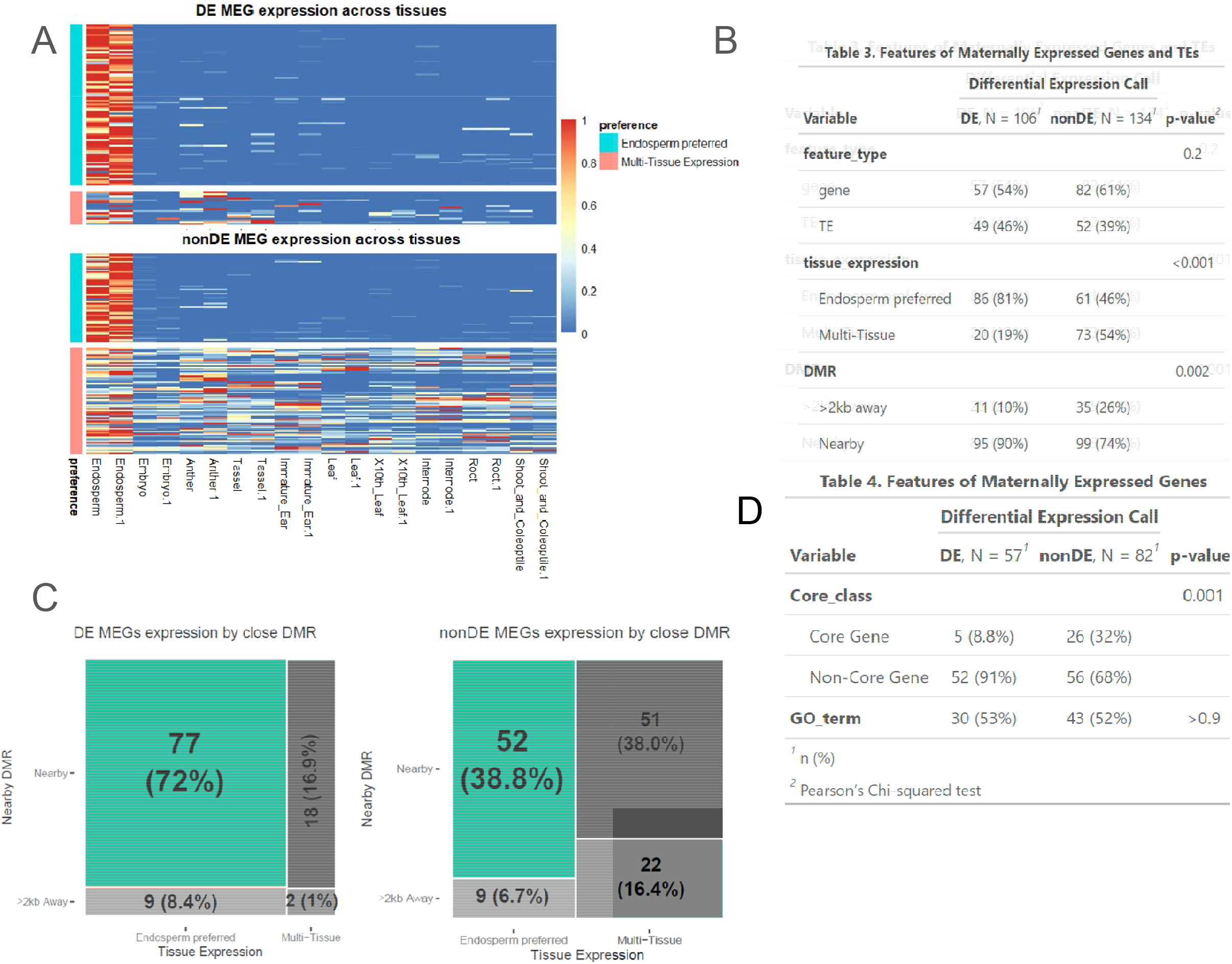
DE MEGs, nonDE MEGs, and conserved imprints. (A) Heatmaps of DE (top) and nonDE (bottom) MEG expression across the W22 expression atlas. Endosperm preferred expression was defined as the sum of the endosperm read samples was greater than the 65% of the sum of all reads, and color scale is reads divided by row sum. This indicates that while both groups display endosperm preferred expression, nonDE MEGs have more variability in the tissues in which they are expressed. This is in line with the prediction that some MEGs are a result of paternal repression and associated with non-endosperm specific expression (B) Table of the features we assessed DE and nonDE MEGs for, as well as p-values. From this we can see that the ratio of TEs to genes is similar in both DE and nonDE MEGs as is the frequency of functional annotations for genes. We can also see that the significant differences between the two groups are nearby DMRs and tissue expression. (C) Endosperm expression preference for DE and nonDE MEGs by distance to closest wt-DMR. 77% of DE MEGs are both endosperm preferred and have a nearby wt-DMR (dark teal) whereas just 38% of nonDE MEGs are both endosperm preferred and have a nearby wt-DMR. (D) Table of the features we assessed between DE and nonDE MEGs limited to genes. To assess core genes, we grouped dispensable genes and genes that do not have an ortholog in B73 into one group called “Non-Core Gene”. We can see that while the number of GO terms isn’t significantly different, the nonDE MEGs are enriched for being core genes.

In addition to differences in DNA methylation and developmental expression patterns, the DE and nonDE MEGs/matTEs differ in several key ways. In contrast to DE MEGs/matTEs, nonDE MEGs/matTEs have a higher proportion of MEGs with 61% of these maternally expressed features being genes, compared to 54% of DE MEGs/matTEs being genes. Additionally, nonDE MEGs had a significantly higher proportion of core genes with 26% to DE MEGs 8.8% (Figure 5 B-D, p-value <0.001). However, the proportion of core genes in nonDE MEGs is intermediate between those values previously seen in all DE vs nonDE genes (Figure 1 C-D).

To further understand the interactions between expression pattern and methylation for MEGs, we assessed CG and CHG methylation at the 2 kb region surrounding the TSS and polyA sites of these genes across mutants and tissues (Figure S4). For DE MEGs, DNA methylation tends to be high in the somatic tissues of leaf and tassel, lower in the endosperm, and returns to near-somatic levels in the *mdr1* and *dng102* mutants. In contrast, nonDE MEGs tend to have low methylation in all tissues surveyed. Interestingly, the nonDE MEGs show an intermediate mean and bimodal distribution of methylation, consistent with a subset of of these genes exhibiting methylation-related de-repression in the endosperm despite having consistent expression in *mdr1* mutant endosperm. These genes may be redundantly regulated with other DNA glycosylases such as DNG102. However, we are unable to assess expression changes in double mutants due to the gametophyte lethality phenotype of double mutants of *mdr1* and *dng102* (Gent *et al*., 2022). Among the nonDE MEGs is *fie1*, a gene which when mutated leads to seed developmental arrest shortly after cellularization (Chaudhury *et al*., 1997). In Arabidopsis, *fie1* is regulated by MDR1’s ortholog, DEMETER, and is a maternally expressed gene. This suggests that although not all maternally de-repressed MEGs lose expression in *mdr1* mutants, the non-DE MEGs that are critical for endosperm development may be redundantly regulated by multiple glycosylases.

## Discussion

MDR1 plays an important role in endosperm development through removal of DNA methylation in the central cell prior to fertilization, establishing a unique epigenetic and transcriptomic environment that is necessary for proper seed development. Though previously targets for demethylation have been identified in *mdr1* mutant endosperm (Gent *et al*., 2022), the changes in transcription were still unknown. Through this study of transcriptional changes in *mdr1* we found 96 genes and 86 TEs that are repressed in *mdr1* and found just 1 gene and 3 TEs that were upregulated (Figure 1C). When assessing the significantly different features of DE and nonDE genes and TEs we found that DE genes/TEs were enriched for endosperm preferred expression, nearby *mdr1*-DMRs, and imprinted expression (Figure 1E). Despite previous studies identifying thousands of *mdr1*-DMRs (Gent *et al*., 2022), there were just 182 genes and TEs displaying differential expression, and these are enriched for having a nearby *mdr1*-DMR when compared to nonDE genes and TEs (Figure 2A). These findings fit with canonical knowledge of the effects of DNA methylation on gene expression, however others have suggested that DNA methylation could impact repressor sequences (Anderson and Springer, 04/2018) and we found little evidence of up-regulated genes in this study.

Additionally we found that among DE features, TEs were enriched when compared to nonDE features (Figure 1E). The association between TEs and DNGs has been previously discussed and identified in endosperm of maize, rice, and Arabidopsis (Hsieh *et al*., 2009; Ibarra *et al*., 2012; Park *et al*., 2017; Gent *et al*., 2022; Yuan *et al*., 2017; Gehring *et al*., 2009). In a previous study of *mdr1* DMRs in maize, the authors found an enrichment for both *mdr1*-DMRs and wt-DMRs overlapping the DHH00002 family of helitrons with approximately 11% of all DMRs overlapping members of this family (Gent *et al*., 2022). We found enrichments for the same DHH00002 family transposons within the DE TEs as well as among the closest TEs to DE genes (Figure 3A/B), which suggests DHH00002 helitrons may have picked up sequence target for MDR1. Further, studies in multiple plant species have found that maternally biased expression is connected with demethylation of nearby transposons (Gehring *et al*., 2009; Hsieh *et al*., 2011; Pignatta *et al*., 2018; Rodrigues *et al*., 2021) suggesting that MDR1 is, in part, responsible for maternally expressed genes through associations with helitron transposons.

One well-established method of imprinting regulation is through removal of methylation on the maternal alleles allowing expression only from the maternal alleles in the endosperm, termed maternal activation (Zhang *et al*., 2014; Gehring *et al*., 2006). Due to MDR1’s activity in the endosperm specifically, we assessed the expression patterns of DE genes to discern whether they were preferentially expressed in the endosperm versus other tissues. We found that generally DE genes were enriched for endosperm preferred expression (Figure 1E) as well as imprinted expression (Figure 4B), which is in line with the activity of MDR1’s Arabidopsis ortholog DEMETER (Gehring *et al*., 2006). Further, our results showing significant overlap between DE genes and previously identified imprinted expression support the notion that MDR1 is a regulator of imprinted expression in maize endosperm, however MDR1 alone does not regulate all imprinted expression. Maize has multiple orthologs (MDR1, DNG102, DNG103, and DNG105) with DEMETER and its three homologs in Arabidopsis. Two of these DNA glycosylases, MDR1 and DNG102, have similar spatiotemporal distributions (Gent *et al*., 2022). In this study we found some overlap in methylation profiles between *mdr1* and *dng102* mutants, suggesting partial redundancy of function between the paralogs (Figure 2D).

Additionally, 106 MEGs lose expression in *mdr1* mutants (Figure 1D). Loss of expression is one way to lose imprinting, establishing MDR1 as a regulator of maternally activated imprinted genes. The 134 MEGs that are not differentially expressed could be explained a couple of ways. First, the nonDE MEGs could be redundantly regulated by MDR1 and DNG102, and thus would only appear nonDE in a double mutant, which is inviable and thus not testable. Alternatively, some nonDE MEGs are likely regulated through a pathway unrelated to MDR1. To discern likely candidates for maternal activation and thus redundant regulation by DNG102 and MDR1, we assessed predictors of maternal activation as defined by (Zhang *et al*., 2014). These predictors included endosperm preferred expression as well as nearby DMRs between embryo and endosperm (wt-DMRs). We found that approximately 39% of nonDE MEGs were both endosperm preferred and have a nearby wt-DMR, indicating that they may be candidates for redundantly regulated genes (Figure 5C). Among this group of nonDE MEGs with features of maternal activation, we found Fie1. Fie1 is a maternally activated gene that, in Arabidopsis, is regulated by DEMETER, MDR1’s ortholog. When mutated, Fie1 leads to seed developmental arrest shortly after cellularization (Chaudhury *et al*., 1997) and is thus essential for endosperm development. Genes with important functions may be redundantly regulated, which could underlie the inviability of an *mdr1/dng102* double mutant. The majority of nonDE maternally activated genes have no known function, however this set of genes may contain additional genes which are strictly regulated and perform important functions in the endosperm. Although there are hypotheses about regulation of MEGs that do not involve maternal demethylation, there is little evidence of how MEGs are regulated otherwise (Batista and Köhler, 2020). The set of MEGs that do not meet any criteria for maternal activation identified in this study could be utilized to investigate other methods of MEG regulation.

In conclusion, a loss of MDR1 in the endosperm leads to decrease of expression for many, but not all, genes with nearby DMRs and a loss of maternal expression for approximately half of the previously identified maternally expressed genes in the endosperm. DE genes share many additional features, including a depletion of GO terms, expression primarily in the endosperm, and close proximity to TEs, particularly helitrons. While this study was limited to single mutants that retain some DNA glycosylase activity from ortholog *DNG102* due to lethality of double mutants, we identified a subset of MEGs that lack any evidence of DNA glycosylase-dependent imprinting, opening the door for future investigation into additional mechanisms of imprinting regulation in maize.

## Materials and Methods

### mdr1 RNA-sequencing and tissue collection

The maize inbred line W22, and a mutant line for the gene *mdr1 (Kermicle, 1995)* in a background of W22 were grown in Athens, GA during the summer of 2021. Homozygous *mdr1* and W22 were self-pollinated, followed by endosperm harvest at 15 days after pollination and manual dissection of approximately 10 kernels from each ear pooled into one biological replicate.

RNA was extracted from each of three biological replicates of W22 and *mdr1* using the Qiagen RNeasy Mini Kit (cat # 74104) after pulverizing wet endosperm in liquid nitrogen. Sample quality checking was performed using an Agilent Bioanalyzer 2100, prior to diluting each sample to 100ng/uL for library preparation and sequencing submission. Paired-end cDNA libraries were prepared using the NEBNext Ultra II Directional RNA kit (cat #E7760S). Samples were then sequenced on the Illumina NovaSeq 6000 using 150 paired-end sequencing at the Iowa State DNA facility, resulting in an average of 300 million reads per sample. Sequence reads were trimmed using cutadapt (Marcel, 2011) (parameters -a AGATCGGAAGAGCACACGTCTGAACTCCAGTCAC -A AGATCGGAAGAGCGTCGTGTAGGGAAAGAGTGTAGATCTCGGTGGTCGCCGTATCATT -m 30 -q 10 --quality-base=33) then aligned to the W22 genome assembly using hisat2 (Kim *et al*., 2019) (parameters -p 6 -k 20). Counts were determined through htseq-count (Anders *et al*., 2014) (parameters -f bam -r pos -s no -t all -i sequence_feature -m union -a 0) using a W22 annotation file the combines gene annotations with TE annotations by first starting with the disjoined TE file, masking exon sequences, then adding full length gene annotations. This approach allows for simultaneous counting of gene and TE expression. The counts were then normalized to reads per million(rpm). Differential expression was determined using R packages, and only features with |>1| Log2FoldChange along with an FDR padj < .05 were called as differentially expressed.

### Features of DE genes analysis

Identification of Core genes defined by conservation within the nested association mapping parental genomes (Hufford *et al*., 2021) was conducted through cross referencing of the pan_gene_matrix_v3_cyverse.csv file, available through cyverse as part of the supplemental data for (Hufford *et al*., 2021), and the pan-zea.v2.pan-genes.tsv file, available at https://download.maizegdb.org/Pan-genes/Pan-zea/, and converting single copy B73v5 gene IDs into single copy W22 gene IDs.

Identification of W22 genes syntenic with sorghum utilized the gene_model_xref_v4c.txt file developed by (Anderson *et al*., 2021) and available at github.com/SNAnderson/Imprinting2020. This file contains W22 gene IDs as well as the syntelogs in sorghum, these columns were compared to DE and nonDE genes IDs from this study to determine synteny. Bedtools closest (Quinlan and Hall, 2010) (parameters -t all -D a) was used to determine distance to DMRs for DE and nonDE genes and TEs, as well as to determine closest TEs to DE and nonDE genes. Visualization was performed in R, all code used to generate Figures is available at https://github.com/kmhiggins/mdr1_DE_analysis.git

### W22 expression atlas read processing

Reads from a W22 expression atlas were downloaded from the NCBI Sequence Read Archive BioProject PRJNA543878 (Monnahan *et al*., 2020). The tissues included in the atlas were: (1) primary root six days after planting, (2) shoot and coleoptile six days after planting, (3) internode at the Vegetative 11 developmental stage, (4) middle of the 10th leaf at the Vegetative 11 developmental stage, (5) middle of the leaf from the ear bearing node at 30□days after pollination, (6) meiotic tassel at the Vegetative 18 developmental stage, (7) immature ear at the Vegetative 18 developmental stage, (8) anthers at the Reproductive 1 developmental stage, (9) endosperm at 16□days after pollination, and (10) embryo at 16□days after pollination. Samples were sequenced on a single run of a HiSeq 2000 as 50 nucleotide single-end reads on an Illumina HiSeq 200 (Illumina, San Diego, CA) at the University of Minnesota BioMedical Genomics Center resulting in an average of 8700000 reads per sample. Reads were then trimmed with cutadapt (parameters -a AGATCGGAAGAGCACACGTCTGAACTCCAGTCAC -m 25 --quality-cutoff=20) (Marcel, 2011) followed by mapping using hisat2(parameters -p 6 -k 20). Count tables were then created using htseq-count (parameters -f bam -r pos -s no -t all -i ID -m union -a 0) (Anders *et al*., 2014) aligned to the same combined gene and TE annotation file described above.

### Hybrid methylome methods

To extract DNA from mature endosperm, seeds were soaked in water for 20 minutes and pericarps, removed with forceps, and embryos with a razor blade. For each biological replicate, three or four endosperms were combined and finely ground with mortars and pestles with liquid N2. DNA was extracted using IBI plant DNA extraction kits (#IB47231). NEBNext^®^ Enzymatic Methyl-seq Kits (#E7120S) used to prepare EM-seq libraries. The input for each library consisted of 200 ng of genomic DNA that had been combined with 1 pg of control pUC19 DNA and 20 pg of control lambda DNA and sonicated to fragments averaging ∼700 bp in length using a Diagenode Bioruptor. The protocol for large insert libraries was followed. Libraries were amplified with 4 PCR cycles. Libraries were sequenced using paired-end 150 nt reads with Illumina HiSeq. Reads were trimmed of adapter sequence using cutadapt, parameters -q 20 -a AGATCGGAAGAGC -A AGATCGGAAGAGC -O. Reads were aligned to the W22 genome to produce bam files and methylation values called in ATCGmap format using BS-Seeker2 (version 2.1.5), parameters -m .03 –aligner=bowtie2 -X 1000 (Guo *et al*., 2013). Alignment files from biological replicates in bam format were combined prior to methylation calling using SAMtools merge (Danecek *et al*., 2021). The CGmapTools version 0.1.2 snv tool was used to identify SNVs from ATCGmap files using default parameters (Guo *et al*., 2018). The asm tool was then used on the resulting vcf files to identify allele-specific methylated sites using default parameters. To increase the speed of this step, the unix split command was used to split each vcf file into a million lines each prior to processing with the asm tool. A custom python script, AlleleSpecificCGmapper.py, was used to extract methylation calls for each genome as separate B73 and W22 CGmap files. The CGmapTools mtr tool was used to calculate average methylation values for each region.

## Acknowledgements

We would like to thank the Iowa State DNA facility for their sequencing services, Conner Valentine for his contribution to RNA extraction, and Dong won Kim for preparation of EM-seq libraries. We additionally want to thank Candice N. Hirsch, Amanda Gilbert, and Yurui Sun for their valuable insights and suggestions. This study was supported in part by resources and technical expertise from the Georgia Advanced Computing Resource Center.

All raw sequencing data generated in this study have been submitted to the NCBI BioProject database (https://www.ncbi.nlm.nih.gov/bioproject/) under accession number PRJNA759188.

## Author contributions

JIG, and SNA designed the research. KMH and JIG generated the data. KMH, JIG, and SNA analyzed data. KMH and SNA wrote the paper with input from all authors.

---

**Table S1:** DE TE Gene Pairs Features.

| DE Gene | DE TE | TE Family | Distance of DMR to Gene | Gene Imprint | TE Imprint | Overlap_Type |
| --- | --- | --- | --- | --- | --- | --- |
| Zm00004b008576 | DHH00002Zm00004b05521 | DHH00002 | <2kb upstream | MEG | MEG | Gene in TE |
| Zm00004b008944 | DHH00002Zm00004b05569 | DHH00002 | >2kb | MEG | no.imprint | Gene in TE |
| Zm00004b019266 | RLG00001Zm00004b16330 | RLG00001 | <2kb upstream | MEG | MEG | Gene in TE |
| Zm00004b020035 | DHH00002Zm00004b01636 | DHH00002 | zero | MEG | MEG | Gene in TE |
| Zm00004b022879 | RLX28628Zm00004b00001 | RLX28628 | zero | MEG | MEG | TE overlaps 5' end of Gene |
| Zm00004b011866 | DHH00002Zm00004b02226 | DHH00002 | zero | MEG | MEG | TE overlaps 3' end of Gene |
| Zm00004b014682 | DHH00002Zm00004b02553 | DHH00002 | >2kb | not imprinted | MEG | TE overlaps 3' end of Gene |
| Zm00004b028628 | DHH00002Zm00004b07508 | DHH00002 | >2kb | MEG | MEG | TE overlaps most of Gene body |
| Zm00004b035625 | DHH02711Zm00004b00065 | DHH02711 | zero | not imprinted | no.imprint | TE overlaps 3' end of Gene |

