## Supplemental Figures for "MDR1 DNA glycosylase regulates the expression of genomically imprinted genes and helitrons"

### Slide 1
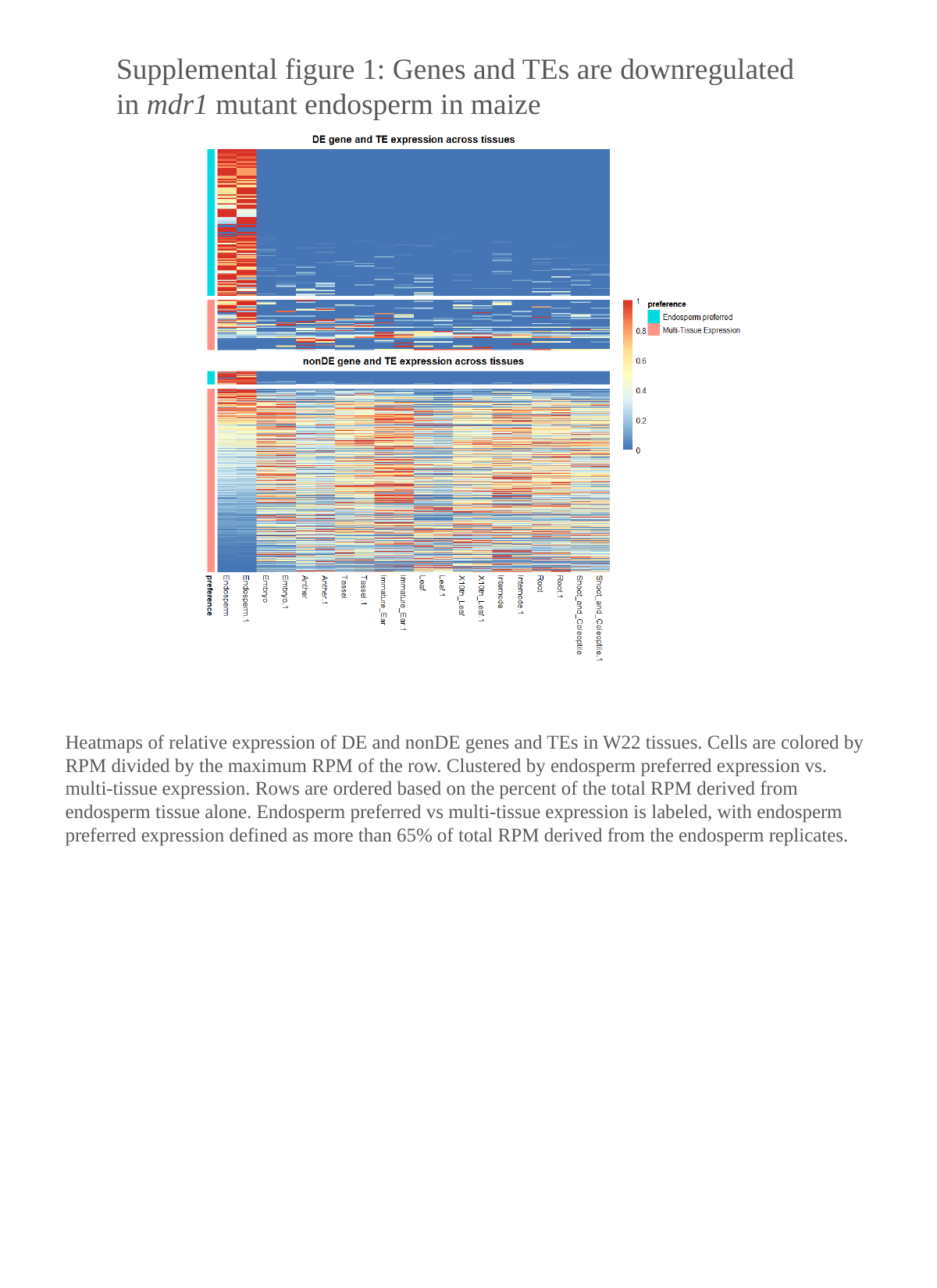

Supplemental figure 1: Genes and TEs are downregulated in mdr1 mutant endosperm in maize
Heatmaps of relative expression of DE and nonDE genes and TEs in W22 tissues. Cells are colored by RPM divided by the maximum RPM of the row. Clustered by endosperm preferred expression vs. multi-tissue expression. Rows are ordered based on the percent of the total RPM derived from endosperm tissue alone. Endosperm preferred vs multi-tissue expression is labeled, with endosperm preferred expression defined as more than 65% of total RPM derived from the endosperm replicates.

### Slide 2
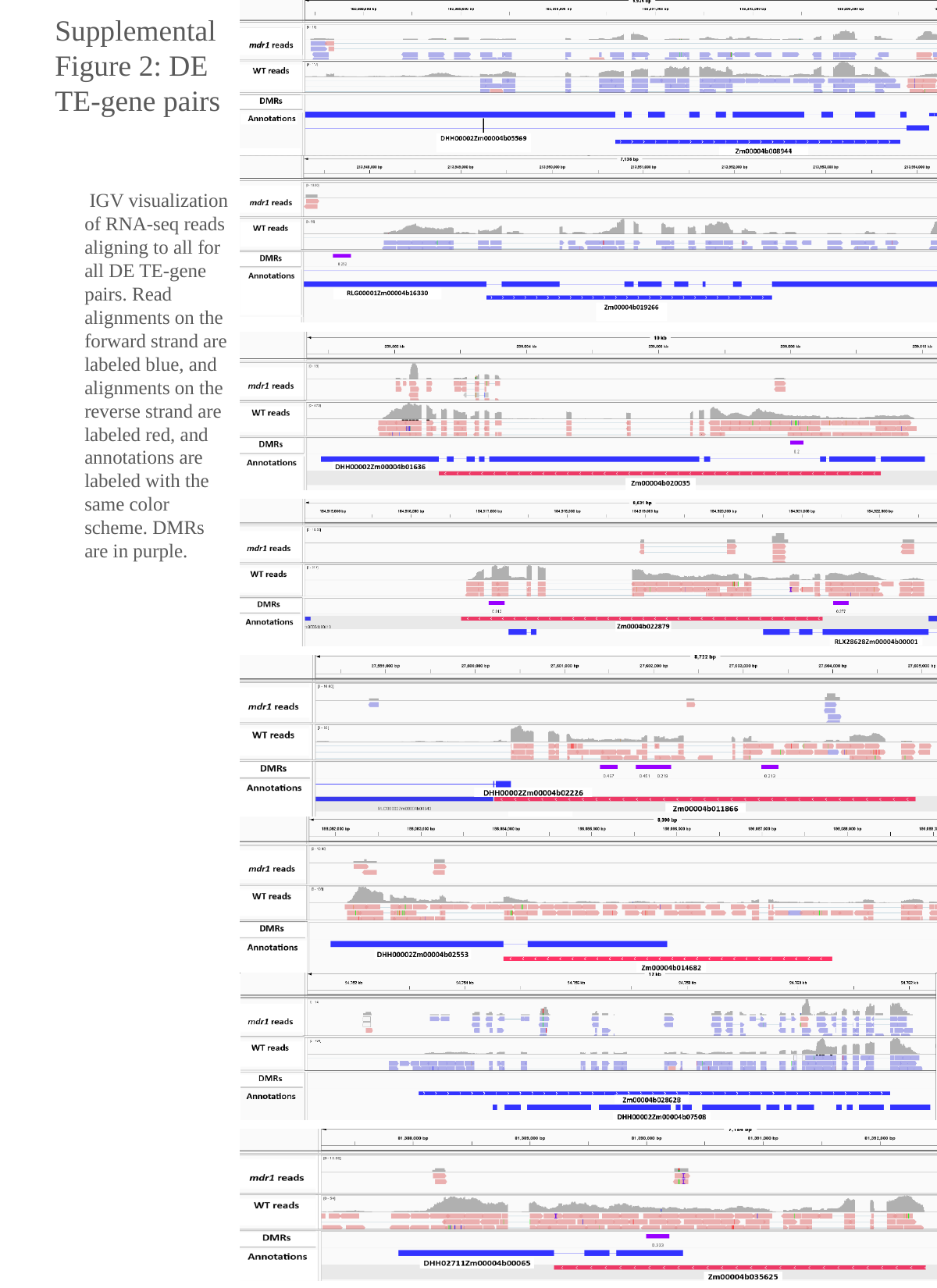

Supplemental Figure 2: DE TE-gene pairs
 IGV visualization of RNA-seq reads aligning to all for all DE TE-gene pairs. Read alignments on the forward strand are labeled blue, and alignments on the reverse strand are labeled red, and annotations are labeled with the same color scheme. DMRs are in purple.

### Slide 3
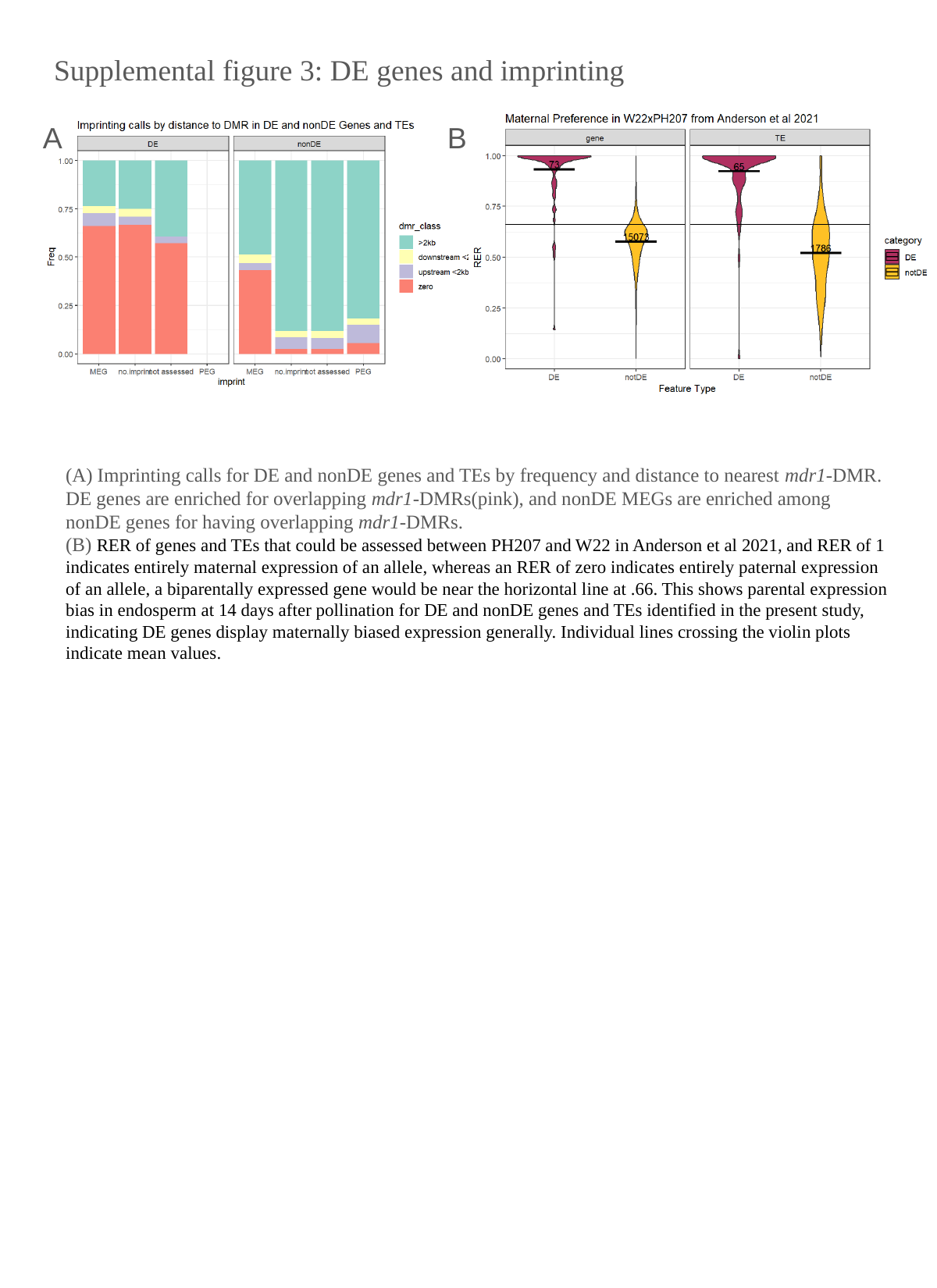

Supplemental figure 3: DE genes and imprinting
A
B
(A) Imprinting calls for DE and nonDE genes and TEs by frequency and distance to nearest mdr1-DMR. DE genes are enriched for overlapping mdr1-DMRs(pink), and nonDE MEGs are enriched among nonDE genes for having overlapping mdr1-DMRs.
(B) RER of genes and TEs that could be assessed between PH207 and W22 in Anderson et al 2021, and RER of 1 indicates entirely maternal expression of an allele, whereas an RER of zero indicates entirely paternal expression of an allele, a biparentally expressed gene would be near the horizontal line at .66. This shows parental expression bias in endosperm at 14 days after pollination for DE and nonDE genes and TEs identified in the present study, indicating DE genes display maternally biased expression generally. Individual lines crossing the violin plots indicate mean values.

### Slide 4
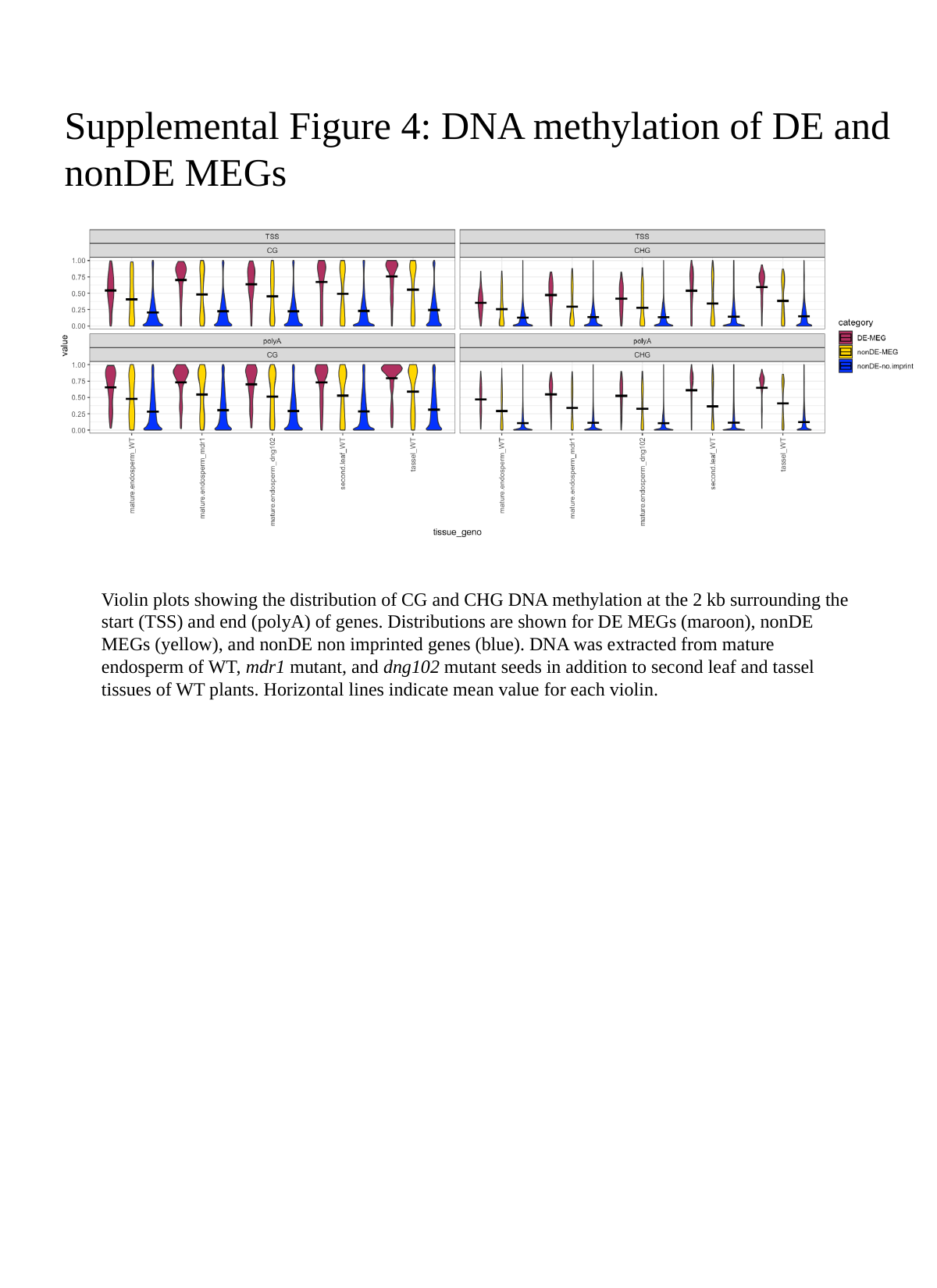

#### Supplemental Figure 4: DNA methylation of DE and nonDE MEGs
Violin plots showing the distribution of CG and CHG DNA methylation at the 2 kb surrounding the start (TSS) and end (polyA) of genes. Distributions are shown for DE MEGs (maroon), nonDE MEGs (yellow), and nonDE non imprinted genes (blue). DNA was extracted from mature endosperm of WT, mdr1 mutant, and dng102 mutant seeds in addition to second leaf and tassel tissues of WT plants. Horizontal lines indicate mean value for each violin.
